# Neural heterogeneity promotes robust learning

**DOI:** 10.1101/2020.12.18.423468

**Authors:** Nicolas Perez-Nieves, Vincent C. H. Leung, Pier Luigi Dragotti, Dan F. M. Goodman

## Abstract

The brain has a hugely diverse, heterogeneous structure. Whether or not heterogeneity at the neural level plays a functional role remains unclear, and has been relatively little explored in models which are often highly homogeneous. We compared the performance of spiking neural networks trained to carry out tasks of real-world difficulty, with varying degrees of heterogeneity, and found that it substantially improved task performance. Learning was more stable and robust, particularly for tasks with a rich temporal structure. In addition, the distribution of neuronal parameters in the trained networks closely matches those observed experimentally. We suggest that the heterogeneity observed in the brain may be more than just the byproduct of noisy processes, but rather may serve an active and important role in allowing animals to learn in changing environments.

**Summary:** Neural heterogeneity is metabolically efficient for learning, and optimal parameter distribution matches experimental data.

## Introduction

The brain is known to be deeply heterogeneous at all scales [1], but it is still not known whether this heterogeneity plays an important functional role or if it is just a byproduct of noisy developmental processes and contingent evolutionary history. A number of hypothetical roles have been suggested (reviewed in [2]), in efficient coding [3–9], reliability [10], working memory [11], and functional specialisation [12]. However, previous studies have largely used simplified tasks or networks, and it remains unknown whether or not heterogeneity can help animals solve complex information processing tasks in natural environments. Recent work has allowed us, for the first time, to train biologically realistic spiking neural networks to carry out these tasks at a high level of performance, using methods derived from machine learning. We used two different learning models [13, 14] to investigate the effect of introducing heterogeneity in the time scales of neurons when performing tasks with realistic and complex temporal structure. We found that it improves overall performance, makes learning more stable and robust, and learns neural parameter distributions that match experimental observations, suggesting that the heterogeneity observed in the brain may be a vital component to its ability to adapt to new environments.

## Results

### Time scale heterogeneity improves learning on tasks with rich temporal structure

We investigated the role of neural heterogeneity in task performance by training recurrent spiking neural networks to classify visual and auditory stimuli with varying degrees of temporal structure. The model used three layers of spiking neurons: an input layer, a recurrently connected layer, and a readout layer used to generate predictions (Figure 1A), a widely used minimal architecture (e.g. Neftci *et al.* [14] and Maass *et al.* [15]). Heterogeneity was introduced by giving each neuron an individual membrane and synaptic time constant. We compared four different conditions: initial values could be either homogeneous or heterogeneous, and training could be either standard or heterogeneous (Figure 1B). Time constants were either initialised with a single value (homogeneous initialisation), or randomly according to a gamma distribution (heterogeneous). The models were trained using surrogate gradient descent [14]. Synaptic weights were always plastic, while time constants were either held fixed at their initial values in the standard training regime, or could be modified in the heterogeneous training regime.

**Figure 1:**
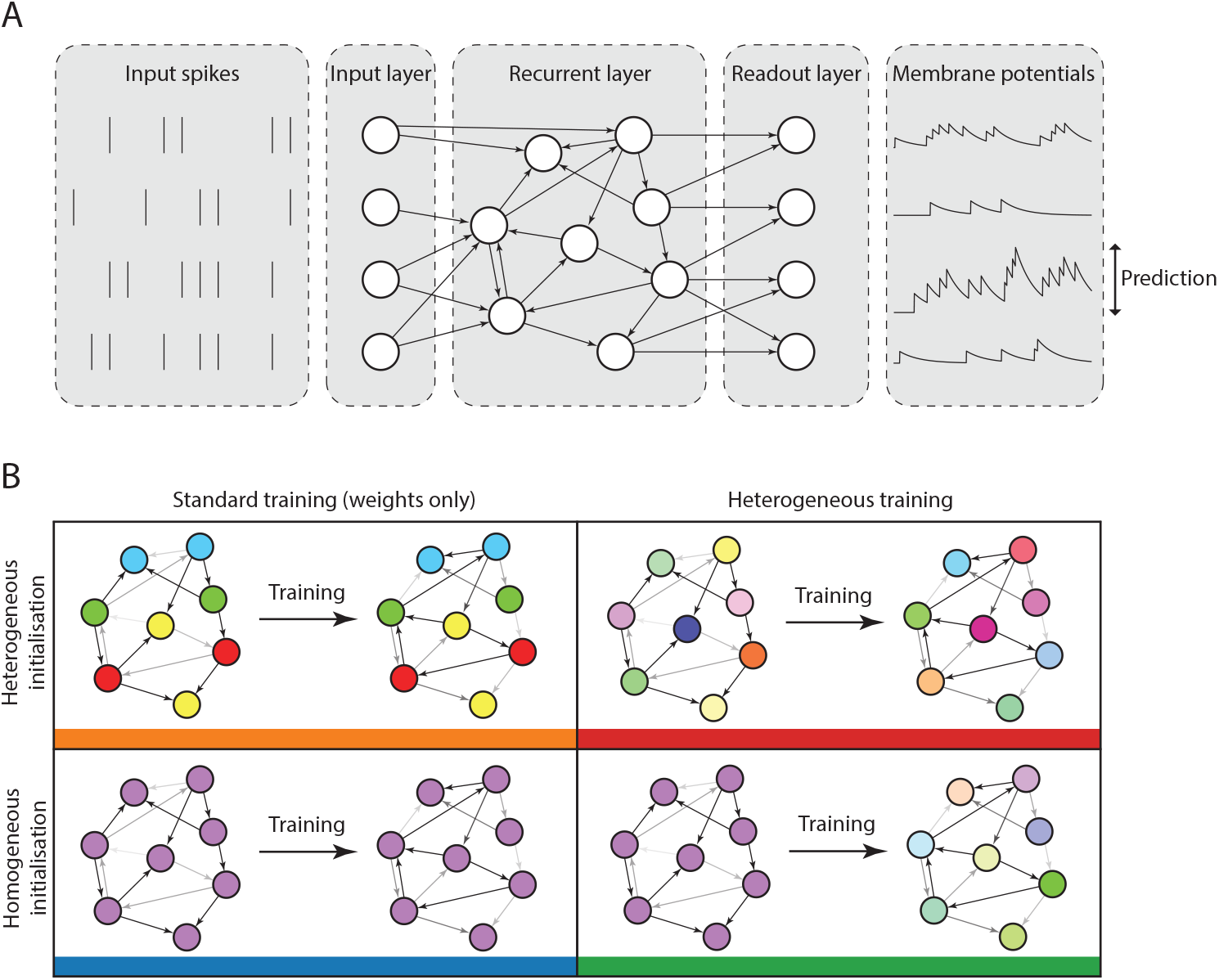
Diagram of network architecture and training configurations. **A.** Model architecture. A layer of input neurons emits spike trains into a recurrently connected layer of spiking neurons which is followed by a readout layer. **B.** Configurations. Training can be either standard (only the synaptic weights are learned) or heterogeneous (the synaptic weights and membrane and synaptic time constants are learned). The initialisation can be homogeneous (all synaptic and membrane time constants are initialised to the same value) or heterogeneous (synaptic and membrane time constants are randomly initialised for each neuron by sampling them from a given probability distribution).

We used five different datasets with varying degrees of temporal structure. Neuromorphic MNIST (N-MNIST; Orchard *et al.* [16]), Fashion-MNIST (F-MNIST; Xiao *et al.* [17], and the DVS128 Gesture dataset [18] feature visual stimuli, while the Spiking Heidelberg Digits (SHD) and Spiking Speech Commands (SSC) datasets [19] are auditory. N-MNIST and DVS128 use a neuromorphic vision sensor to generate spiking activity, by moving the sensor with a static visual image of handwritten digits (N-MNIST) or by recording humans making hand gestures (DVS128). F-MNIST is a dataset of static images that is widely used in machine learning, which we converted into spike times by treating the image intensities as input currents to model neurons, so that higher intensity pixels would lead to earlier spikes, and lower intensity to later spikes. Both SHD and SSC use a detailed model of the activity of bushy cells in the cochlear nucleus, in response to spoken digits (SHD) or commands (SSC). Of these datasets, N-MNIST and F-MNIST have minimal temporal structure, as they are generated from static images. DVS128 has some temporal structure as it is recorded motion, but it is possible to perform well at this task by discarding the temporal information. The auditory tasks SHD and SSC by contrast have very rich temporal structure.

We found that heterogeneity in time constants had a profound impact on performance on those training datasets where information was encoded in the precise timing of input spikes (Table 1, Figure 2A). On the most temporally complex auditory tasks, accuracy improved by a factor of around 15-20%, while for the least temporally complex task N-MNIST we saw no improvement at all. For the gesture dataset DVS128 we can identify the source of the (intermediate) improvement as the heterogeneous models being better able to distinguish between spatially similar but temporally different gestures, such as clockwise and anticlockwise versions of the same gesture (See Supp. Figure 10A-B). This suggests that we might see greater improvements for a richer version of this dataset in which temporal structure was more important.

**Table 1:** Testing accuracy percentage over different datasets and training methods. Effect of initialisation and training configuration on performance, on datasets of increasing temporal complexity. Initialisation can be homogeneous (all time constants the same) or heterogeneous (random initialisation), and training can be standard (only synaptic weights learned) or heterogeneous (time constants can also be learned). N-MNIST and F-MNIST are static image datasets with little temporal structure, DVS128 is video gestures, and SHD and SSC are temporally complex auditory datasets.

| Initialisation | Training | N-MNIST | F-MNIST | DVS128 | SHD | SSC |
| --- | --- | --- | --- | --- | --- | --- |
| Homog. | Standard | 97.4±0.0 | 80.1±7.4 | 76.9±0.8 | 71.7±1.0 | 49.7±0.4 |
| Heterog. | Standard | 97.5±0.0 | 87.9±0.1 | 79.5±1.0 | 73.7±1.1 | 53.7±0.7 |
| Homog. | Heterog. | 96.6±0.2 | 79.7±7.4 | 81.2±0.8 | 82.7±0.8 | 56.6±0.7 |
| Heterog. | Heterog. | 97.3±0.1 | 87.5±0.1 | 82.1±0.8 | 81.7±0.8 | 60.1±0.7 |
| Chance level |  | 10.0 | 10.0 | 10.0 | 5.0 | 2.9 |

**Figure 2:**
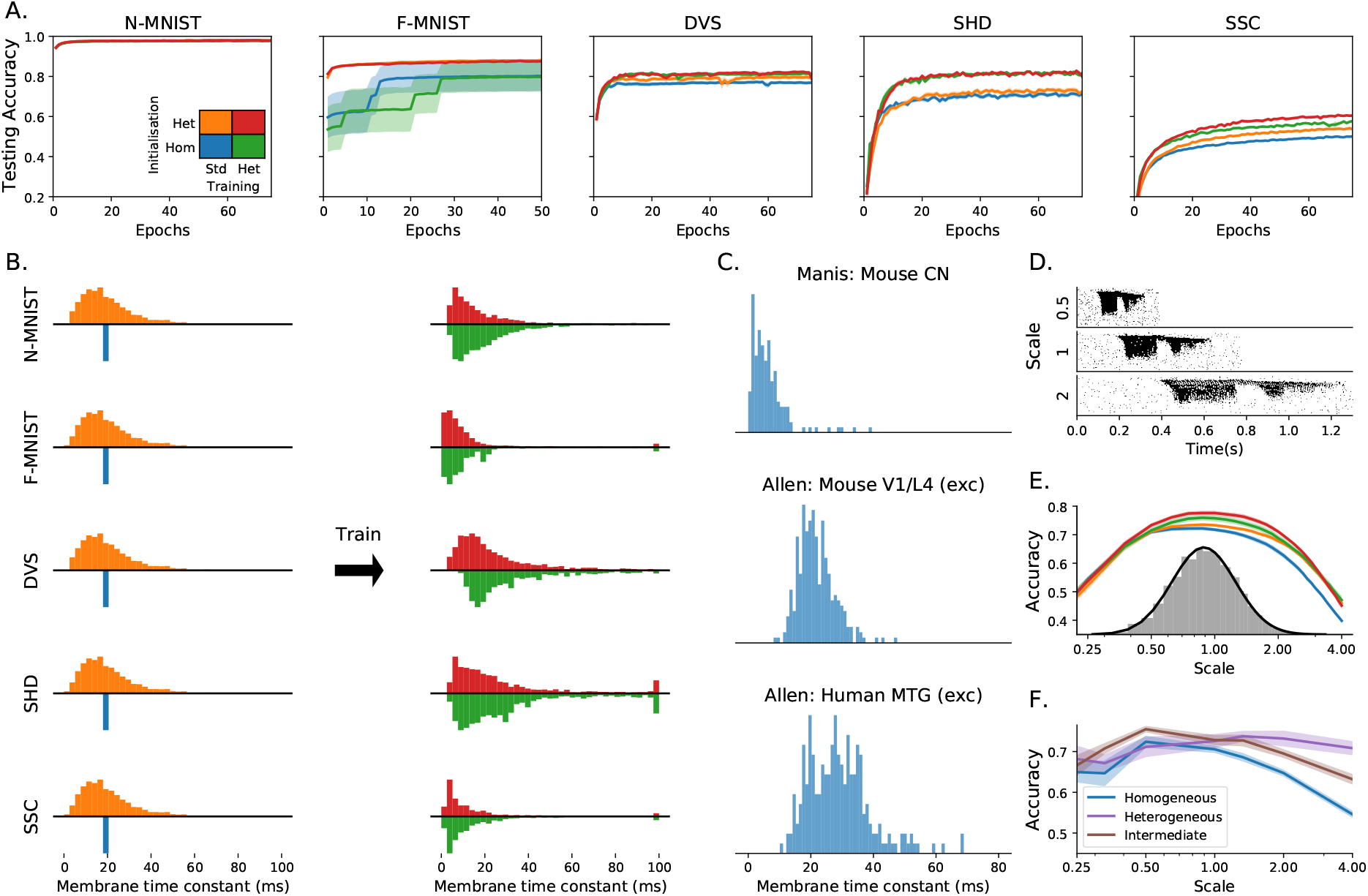
Impact of training configuration and temporal structure of the dataset on the testing accuracy, membrane time constant distributions and performance when training at different time scales. **A.** Improvements in accuracy in testing data, for datasets with temporal complexity low (N-MNIST, F-MNIST), intermediate (DVS) and high (SHD). Shaded areas correspond to standard error in the mean over 10 trials. Initialisation can be homogeneous (blue/green) or heterogeneous (orange/red), and training can be standard, weights only (blue/orange) or heterogeneous including time constants (green/red). Heterogeneous configurations achieve a better test accuracy on the more temporally complex datasets. Heterogeneous initialisation also results in a more stable and robust training trajectory for F-MNIST, leading to better performance overall. **B.** Membrane time constant distributions before (left) and after (right) training for each dataset. Histograms above the axis represent heterogeneous initialisation, and below the axis homogeneous initialisation. In the case of standard training (weights only), the initial distribution (left) is the same as the final distribution of time constants after training. **C.** Experimentally observed distributions of time constants for (top to bottom): mouse cochlear nucleus, multiple cell types (172 cells); mouse V1 layer 4, spiny (putatively excitatory) cells (164 cells); human middle temporal gyrus, spiny cells (236 cells). **D.** Raster plot on input spikes from a single sample of the SHD dataset (spoken digits) at three different time scales. **E.** Accuracy on the SHD dataset after training on a variety of time scales (randomly selected from the grey distribution) for the four configurations described in (A). **F.** Accuracy on the SHD dataset when the initial distribution of time constants is tuned for time scale 1.0, but the training and testing is done at different time scales.

We verified that our results were due to heterogeneity and not simply to a better tuning of time constants in two ways. Firstly, we performed a grid search across all homogeneous time constants for the SHD dataset and used the best values for our comparison. Secondly, we observe that the distribution of time constants after training is very similar and heterogeneous regardless of whether it was initialised with a homogeneous or heterogeneous distribution (Figure 2B), indicating that the heterogeneous distribution is optimal.

Introducing heterogeneity allows for a large increase in performance at the cost of only a very small increase in the number of parameters (0.23% for SHD, because the vast majority of parameters are synaptic weights), and without using any additional neurons or synapses. Heterogeneity is therefore a metabolically efficient strategy. It is also a computationally efficient strategy of interest to neuromorphic computing, because adding heterogeneity adds *O*(*n*) to memory use and computation time, while adding more neurons adds *O*(*n*^2^). Further, in some neuromorphic systems like BrainScaleS this heterogeneity is already present as part of the manufacturing process [20].

Note that it is possible to obtain better performance using a larger number of neurons. For example, Neftci *et al.* [14] obtained a performance of 83.2% on the SHD dataset without heterogeneity using 1024 neurons and data augmentation techniques, whereas we obtained 82.7% using 128 neurons and no data augmentation. We focus on smaller networks here for two reasons. Firstly, we wanted to systematically investigate the effect of different training regimes, and current limitations of surrogate gradient descent mean that each training session takes several days. Secondly, with larger numbers of neurons, performance even without heterogeneity approaches the ceiling on these tasks (which are still simple in comparison to those faced by animals in real environments), making it more difficult to see the effect of different architectures. Even with this limitation to small networks, heterogeneity confers such an advantage that our results for the SSC dataset are state of the art (for spiking neural networks) by a large margin.

We also tested the effect of introducing heterogeneity of other neuron parameters, such as the firing threshold and reset potential, but found that it made no appreciable difference. This was because for our model, changing these is almost equivalent to a simple scaling of the membrane potential variable. By contrast, Bellec *et al.* [21] found that introducing an adaptive threshold did improve performance, presumably because it allows for much richer temporal dynamics.

### Predicted time constant distributions match experimental data

In all our tasks, the distribution of time constants after training approximately but not exactly fit a log normal or gamma distribution (with different parameters for each task), and are consistent across different training runs (Supplementary Materials Figure 4 and Figure 5), suggesting that the learned distributions may be optimal. Using publicly available datasets including time constants recorded in large numbers of neurons in different animals and brain regions [22–25], we found very similar distributions to those we predicted (Figure 2C). The parameters for these distributions are different for each animal and region, just as for different tasks in our simulations. Interestingly, the distribution parameters are also different for each cell type in the experimental data, a feature not replicated in our simulations as all cells are identical. This suggests that introducing further diversity in terms of different cell types may lead to even better performance.

### Heterogeneity improves speech learning across time scales

Sensory signals such as speech and motion can be recognised across a range of speeds. We tested the role of heterogeneity in learning a circuit that can function across a wide range of speeds. We augmented the SHD spoken digits datasets to include faster or slower versions of the samples, multiplying all spike times by a temporal scale as an extremely simplified model that captures a part of the difficulty of this task (Figure 2D). During training, temporal scales were randomly selected from a distribution roughly matching human syllabic rate distributions [26]. The heterogeneous configurations performed as well or better at all time scales (Figure 2E), and in particular were able to generalise better to time scales outside the training distribution (e.g. accuracy of 47% close to a time scale of 4, around 7% higher than the homogeneous network, where chance performance would be 5%). Heterogeneous initialisation alone was sufficient to achieve this better generalisation performance, while fine tuning the distribution of time constants with heterogeneous training improved the peak performance but gave no additional ability to generalise.

### Heterogeneity improves robustness against mistuned learning

We tested the hypothesis that heterogeneity can provide robustness with two experiments where the *hyperparameters* were mistuned, that is where the initial distributions and learning parameters were chosen to give the best performance for one distribution, but the actual training and testing is done on a different distribution.

In the first experiment, we took the augmented SHD spoken digits dataset from the previous section, selected the hyperparameters to give the best performance at a time scale of 1, but then trained and tested the network at a different time scale (Figure 2F). We used training of weights only, since allowing retraining of time constants lets the network cancel out the effect of changing the time scale. With a homogeneous or narrow heterogeneous initial distribution, performance falls off for time scales far from the optimal one, particularly for larger time scales (slower stimuli). However, a wide heterogeneous initial distribution allows for good performance across all time scales tested, at the cost of slightly lower peak performance at the best time scale. We tested whether or not this was solely due to the presence of a few neurons with long time constants by introducing an intermediate distribution where the majority of time constants were the same as the homogeneous case, with a small minority of much longer time constants. The performance of this distribution was intermediate between the homogeneous and heterogeneous distributions, a pattern that is repeated in our second experiment below.

In the second experiment, we switched to a very different learning paradigm, FORCE training of spiking neural networks [13] to replay a learned signal, in this case a recording of a zebra finch call (Figure 3A; from Blättler & Hahnloser [27]). This method does not allow for heterogeneous training (tuning time constants), so we only tested the role of untrained heterogeneous neuron parameters. We tested three configurations: fully homogeneous (single 20ms time constant as in the original paper); intermediate (each neuron randomly assigned a fixed fast 20ms or slow 100ms time constant); or fully heterogeneous (each neuron randomly assigned a time constant drawn from a gamma distribution).

**Figure 3:**
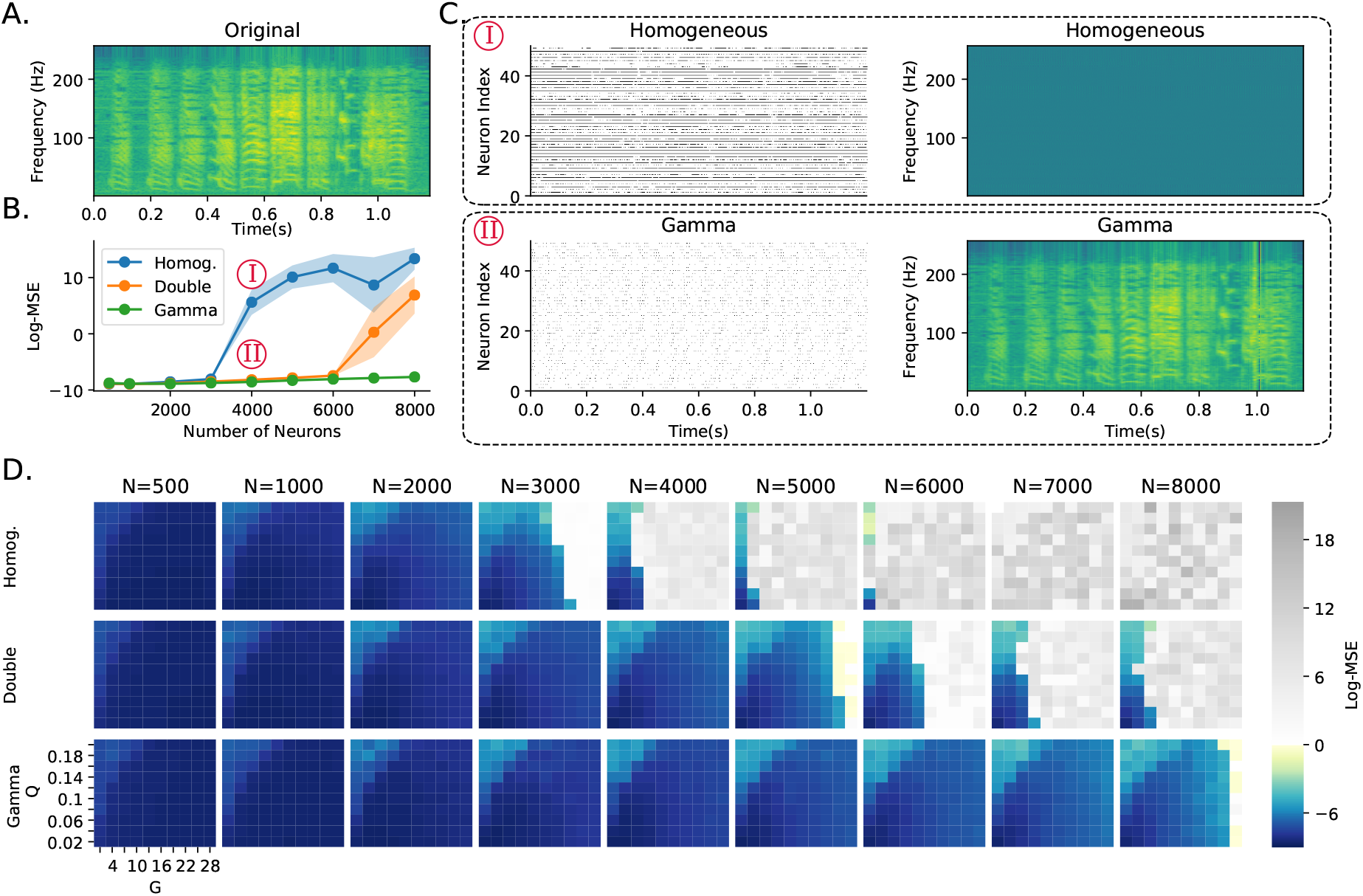
Robustness to learning hyperparameter mistuning. **A.** Spectrogram of a zebra finch. The network has to learn to reproduce this spectrogram, chosen for its spectrotemporal complexity. **B** Error for three networks at different network sizes (hyperparameters were chosen to optimise performance at *N* = 1000 neurons). Networks are fully homogeneous (*Homog*); intermediate, where each neuron is randomly assigned slow or fast dynamics (*Double*); or fully heterogeneous, where each neuron has a random time constant drawn from a gamma distribution (*Gamma*). **C**. Raster plots of 50 neurons randomly chosen, and reconstructed spectrograms under fully homogeneous and fully heterogeneous (Gamma) conditions for *N* = 4000 neurons as indicated in (B). **D**. Reconstruction error. Each row is one of the conditions in (B). Each column is a network size. The axes of each image give the learning hyperparameters (*G* and *Q*). Grey pixels correspond to log mean square error above 0, corresponding to a complete failure to reconstruct the spectrogram. The larger the coloured region, the more robust the network is, and the less tuning is required.

Nicola & Clopath [13] showed that network performance is highly dependent on two hyperparameters (*G* and *Q* in their paper). We therefore tuned these hyperparameters for a network of a fixed size (*N* = 1000 neurons) and ran the training and testing for networks of different sizes (Figure 3B). As the network size started to diverge, the homogeneous network began to make large errors, while the fully heterogeneous network was able to give low errors for all network sizes. The intermediate network was able to function well across a wider range of network sizes than the fully homogeneous network, but still eventually failed for the largest network sizes. At these large network sizes, the homogeneous network becomes saturated, leading to poor performance (Figure 3C). The robustness of the heterogeneous version of the network can be measured by the area of the hyperparameter space that leads to good performance (Figure 3D). Adding partial or full heterogeneity leads to an improvement in learning for all points in the hyperparameter space. again suggesting that it can improve robustness of learning in a wide range of situations.

## Discussion

We trained spiking neural networks at difficult classification tasks, either forcing all time constants to be the same (homogeneous) or allowing them to be different (heterogeneous). We found that introducing heterogeneity improved overall performance across a range of tasks and training methods, but particularly so on tasks with richer intrinsic temporal structure. Learning was more robust, in that the networks were able to learn across a range of different environments, and when the hyperparameters of learning were mistuned. When the learning rule was allowed to tune the time constants as well as synaptic weights, a consistent distribution of time constants was found, akin to a log normal or gamma distribution, and this qualitatively matched time constants measured in experimental data. Note that we do not claim that the nervous system tunes time constants during its lifetime to optimise task performance, we only use our methods to find the optimal distribution of time constants.

We conclude from this that neural heterogeneity is a metabolically efficient strategy for the brain. Heterogeneous networks have no additional cost in terms of the number of neurons or synapses, and perform as well as homogeneous networks which have an order of magnitude more neurons. This gain also extends to neuromorphic computing systems, as adding heterogeneity to the neuron model adds an additional time and memory cost of only *O*(*n*), while adding more neurons has a cost of *O*(*n*^2^). In addition to their overall performance being better, heterogeneous networks are more robust and able to learn across a wider range of environments, which is clearly ethologically advantageous. Again, this has a corresponding benefit to neuromorphic computing and potentially machine learning more generally, in that it reduces the cost of hyperparameter tuning, which is often one of the largest costs for developing these models.

The question remains as to the extent of time constant tuning in real nervous systems. It could be the case that the heterogeneous distribution of time constants observed in different animals and brain regions (Figure 2C) is simply a product of noisy developmental processes. This may be so, but our results show that these distributions closely match the optimal ones found by simulation which confer a substantial computational advantage, and it therefore seems likely that the brain makes use of this advantage. We found that any degree of heterogeneity improves performance, but that the best performance could be found by tuning the distribution of time constants to match the task. Without a more detailed model of these specific brain regions and the tasks they solve, it is difficult to conclude whether or not the precise distributions observed are tuned to those tasks or not, and indeed having a less precisely tuned distribution may lead to greater robustness in uncertain environments.

A number of studies have used heterogeneous or tunable time constants [28–30], but these have generally been focussed on maximising performance for neuromorphic applications, and not considering the potential role in real nervous systems. In particular, we have shown that: heterogeneity is particularly important for the type of temporally complex tasks faced in real environments, as compared to the static ones often considered in machine learning; heterogeneity confers robustness allowing for learning in a wide range of environments; optimal distributions of time constants are consistent across training runs and match experimental data; and that our results are not specific to a particular task or training method.

The methods used here are very computationally demanding, and this has limited us to investigating very small networks (hundreds of neurons). Indeed, we estimate that in the preparation of this paper we used approximately 2 years of GPU computing. Finding new algorithms to allow us to scale these methods to larger networks will be a critical task for the field.

Beyond this, it would be interesting to see to what extent different forms of heterogeneity confer other advantages, such as spatial heterogeneity as well as temporal. We observed that in the brain, different cell types have different stereotyped distributions of time constants, and it would be interesting to extend our methods to networks with multiple cell types, including more biophysically detailed cell models.

Our computational results show a compelling advantage for heterogeneity, and this makes intuitive sense. Having heterogeneous time constants in a layer allows the network to integrate incoming spikes at different time scales, corresponding to shorter or longer memory trace, thus allowing the readout layer to capture information at several scales and represent a richer set of functions. It would be very valuable to extend this line of thought and find a rigorous theoretical explanation of the advantage of heterogeneity.

## Data and coda availability statement

The code for this work and links to the different datasets used is available at https://github.com/npvoid/neural_heterogeneity.

## Acknowledgments

We are immensely grateful to the Allen Institute and Paul Manis for publicly sharing their databases that allowed us to estimate time constant distributions in the brain. Releasing this data is not only immensely generous and essential for this work, but more generally it accelerates the pace of science and represents an optimistic vision of the future.

## Funding

The support of the EPSRC Centre for Doctoral Training in High Performance Embedded and Distributed Systems (HiPEDS, Grant Reference EP/L016796/1) and the Imperial College President’s PhD Scholarship is gratefully acknowledged.

## Author contributions

N.P.-N.: Conceptualisation, Methodology, Software, Validation, Visualisation, Writing - Original Draft; V.C.H.L.: Methodology, Software, Validation, Visualisation, Writing - Review & Editing; P.L.D.: Writing - Review & Editing, Supervision, Project administration; D.F.M.G.: Conceptualisation, Resources, Data Curation, Writing - Review & Editing, Supervision, Project administration.

**List of Supplementary Materials**

- Materials and Methods
- Table S1
- Figures S2-S8

## Supplementary materials

### Materials and Methods

#### Neuron and synaptic models

We use the Leaky Integrate and Fire (LIF) neuron model in all our simulations. In this model the membrane potential of the *i*-th neuron in the *l*-th layer 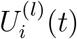 varies over time following (1).

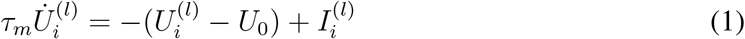

Here, *τ_m_* is the membrane time constant, *U*_0_ is the resting potential and 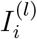 is the input current. When the membrane potential reaches the threshold value *U_th_* a spike is emitted, *U_i_*(*t*) resets to the reset potential *U_r_* and then enters a refractory period that lasts *t_ref_* seconds where the neuron cannot spike.

Spikes emitted by the *j*-th neuron in layer *l*−1 at a finite set of times 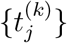 can be formalised as a spike train 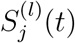 defined as in (2)

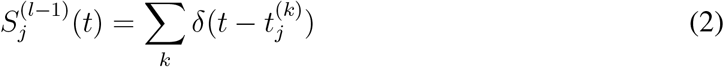

The input current 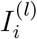 is obtained from the spike trains of all presynaptic neurons *j* connected to neuron *i* following (3)

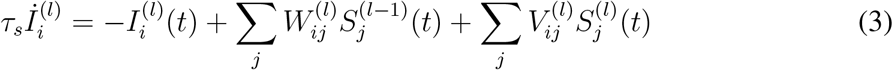

Here *τ_s_* is the synaptic time constant, 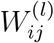 is the feed-forward synaptic weight from neuron *j* in layer *l*−1 to neuron *i* in layer *l* and 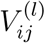 is the recurrent weight from neuron *j* in layer *l* to neuron *i* in layer *l*.

Thus, a LIF neuron is fully defined by six parameters *τ_m_, τ_s_, U_th_, U*_0_*, U_r_, t_ref_* plus its synaptic weights 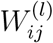 and 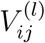. We refer to these as the **neuron parameters** and **weights** respectively.

Since we are considering the cases where these parameters may be different for each neuron in the population we should actually refer to *τ_m,i_*, *τ_s,i_*, *U_th,i_*, *U*_0*,i*_,*U_r,i_*, *t_ref,i_*. However, for notational simplicity we will drop the *i* subscript and it will be assumed that these parameters can be different for each neuron in a population.

#### Neural and synaptic model discretisation

In order to implement the LIF model in a computer it is necessary to discretise it. Assuming a very small simulation time step Δ*t*, (3) can be discretised to

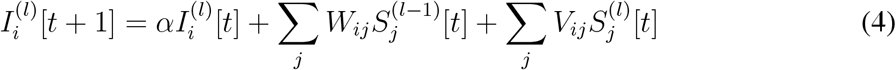

With *α* = exp(−Δ*t/τ_s_*). Similarly, (1) becomes

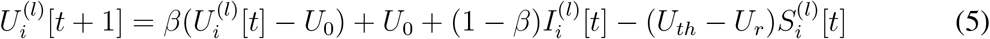

With *β* = exp(−Δ*t/τ_m_*). Finally, the spiking mechanism

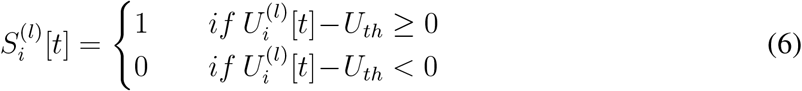

Notice how the last term in (5) introduces the membrane potential resetting. This would only work if we assume that the neuron potential at spiking time was exactly equal to *U_th_*. This may not necessarily be the case since membrane potential update that crossed the threshold may result in 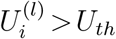 and then the resetting mechanism will not set the membrane potential to *U_r_*. However, we found that this has a negligible effect in our simulations.

#### Surrogate Gradient Descent Training

With the discretisation introduced in the previous section, a spiking layer consists of three cascaded sub-layers: current (4), membrane (5) and spike (6). The current and membrane sublayers have access to its previous state and thus, they can be seen as a particular case of recurrent neural network (RNN). Note that while each neuron is a recurrent unit since it has access to its own previous state, different neurons in the same spiking layer will only be connected if any of the non-diagonal elements of ***V***^(*l*)^ is non-zero. In other words, all SNNs built using this model are RNNs but not all SNNs are RSNNs.

We can cascade *L* spiking layers to conform a Deep Spiking Neural Network analogous to a conventional Deep Neural Network and train it using gradient descent. However, since equation (6) is non-differentiable, we need to modify the backwards pass as in [14] so that BPTT algorithm can be used to update the network parameters.

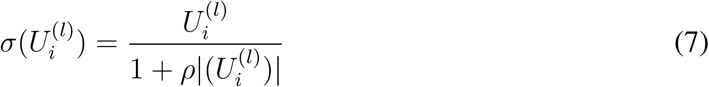

This means that while in the forward pass the network follows a step function as in (6), in the backwards pass it follows a sigmoidal function (7), with steepness set by *ρ*.

We can now use gradient descent to optimise the synaptic weights ***W***^(*l*)^ and ***V***^(*l*)^ as in conventional deep learning. We can also optimise the spiking neuron specific parameters *U_th_, U*_0_*, U_r_* since they can be seen as bias terms. The time constants can also be indirectly optimised by training *α* and *β* which can be seen as forgetting factors.

We apply a clipping function to *α* and *β* after every update.

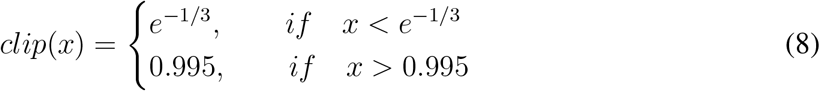

In order to ensure stability, the forgetting factors have to be less than 1. Otherwise, the current and membrane potential would grow exponentially. Secondly to make the system causal these factors cannot be less than zero. This however, would allow for arbitrarily small time constants which would not have any meaning given a finite time resolution Δ*t*. Thus, we constrain the time constants to be at least 3Δ*t*. We also set clipping limits for *U_th_, U*_0_*, U_r_* such that they are always between the ranges specified in Table 2.

**Table 2:** Parameter initialisation for the different configurations

| Parameter | HomInit | HetInit |
| --- | --- | --- |
| $\tau_m$ | $\bar{\tau}_m$ | $\Gamma(3, \bar{\tau}_m/3)$ |
| $\tau_s$ | $\bar{\tau}_s$ | $\Gamma(3, \bar{\tau}_s/3)$ |
| $U_{th}$ | $U_{th}$ | $\mathcal{U}(0.5, 1.5)$ |
| $U_0$ | $U_0$ | $\mathcal{U}(-0.5, 0.5)$ |
| $U_r$ | $U_r$ | $\mathcal{U}(-0.5, 0.5)$ |

There are several ways in which the neuron parameters may be trained. One possibility is to make all neurons in a layer share the same neuron parameters. That is, a single value of *U_th_, U*_0_*, U_r_, α, β* is trained and shared by all neurons in a layer. Another possibility is to optimise each of these parameters in each neuron individually as we have done in our experiments. We also always trained the weight matrices ***W***^(*l*))^ and ***V***^(*l*))^. Training was done by using automatic differentiation on PyTorch ([31]) and Adam optimiser with learning rate 10^−3^ and betas (0.9, 0.999).

In all surrogate gradient descent experiments a single recurrent spiking layer with 128 neurons received all input spikes. This recurrent layer is followed by a feedforward readout layer with *U_th_* set to infinity and with as many neurons as classes in the dataset.

For the loss function, we follow the *max-over-time* loss in [19] to take the maximal membrane potential over the entire time in the readout layer. We then take these potentials and compute the cross-entropy loss

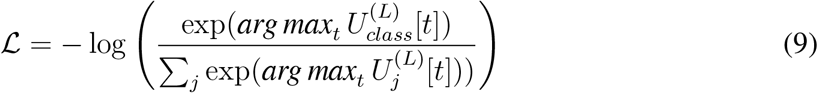

where *class* corresponds to the readout neuron index of the correct label for a given sample. The loss is computed as the average of *N_batch_* training samples. This was repeated for a total of *N_epochs_*.

In order to improve generalisation we added noise to the input by adding spikes following a Poisson process with rate 1.2 Hz and deleting spikes with probability 0.001.

The parameters used for the network are given in Table 2 and Table 3 unless otherwise specified. In Figure 2F we used a log-normal distribution in which we ensured the mode was the same as in the Gamma distribution we used for the other experiments (see Table 2) but we scaled the standard deviation to be *f* times that of the original Gamma distribution. We used *f* = 4 for the heterogeneous case. For the intermediate case all neurons were initialised as in the Homogeneous configuration but 5% of them selected randomly were given the largest time constant value allowed of 100ms.

**Table 3:**
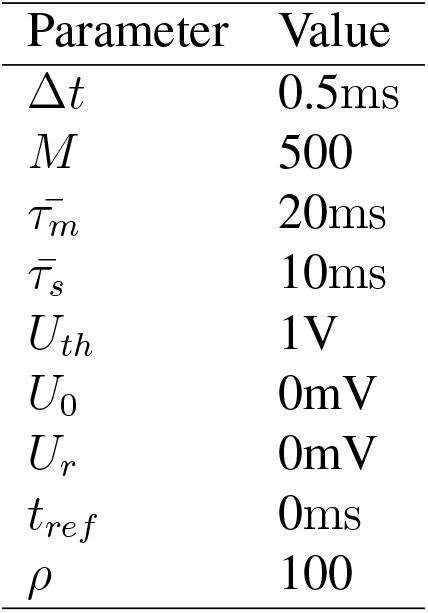
FORCE network parameters

| Parameter | Value |
| --- | --- |
| $\Delta t$ | 0.5ms |
| $M$ | 500 |
| $\bar{\tau}_m$ | 20ms |
| $\bar{\tau}_s$ | 10ms |
| $U_{th}$ | 1V |
| $U_0$ | 0mV |
| $U_r$ | 0mV |
| $t_{ref}$ | 0ms |
| $\rho$ | 100 |

All states *I_i_*(*l*)[0] and *U_i_*(*l*)[0] are initialised to 0. For the weights ***W*** and ***V***, we independently sampled from a uniform distribution 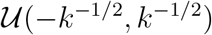, with *k* being the number of afferent connections ([32]).

#### FORCE Training

The FORCE method is used to train a network consisting of a single recurrent layer of LIF neurons as in [13]. In this method, there are no feedforward weights and only the recurrent weights ***V*** are trained. We can express these weights as

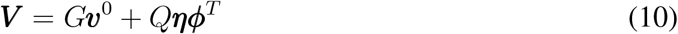

The first term in (10), namely *G****v***^0^, remains static during training and it is initialised to set the network into chaotic spiking. The learned component of the weights ***ϕ***^*T*^ ∈ℝ^*K*×*N*^ is updated using the Recursive Least Squares algorithm. The vector ***η*** ∈ ℝ^*N*×*K*^ serves as a decoder and it is static during learning. The constants *G* and *Q* govern the ratio between chaotic and learned weights.

With this definition of ***V*** we can write the currents into the neurons as the sum ***I***[*t*] = ***I***_*G*_[*t*]+***I***_*Q*_[*t*] (we dropped the layer *l* superscript since we only have a single layer) where we define

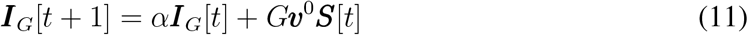

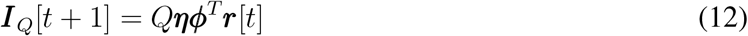

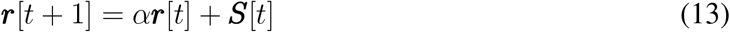

In order to stabilise the network dynamics we add a High Dimensional Temporal Signal (HDTS) as in [13]. This is an *M* dimensional periodic signal ***z***[*t*]. Given the HDTS period *T* , we split the interval [0, *T*] into *M* subintervals *I_m_, m* = 1, … , *M* such that each of the components of ***z***[*t*] is given by

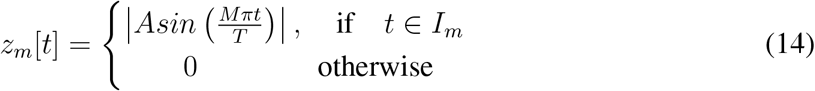

This signal is then projected onto the neurons leaving equation (4) as

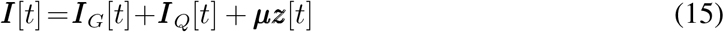

where vector ***μ*** ∈ ℝ^*N*×*M*^ is just a decoder similar to ***η*** in (10)

The aim of FORCE learning is to approximate a *K*-dimensional time varying teaching signal ***x***[*t*]. The vector ***r***[*t*] is used to obtain an approximant of the desired signal

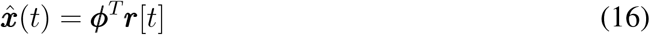

The weights are updated using the RLS learning rule according to:

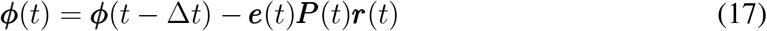

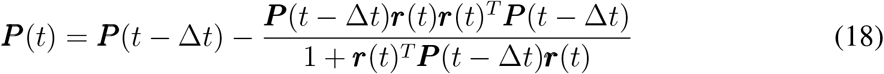

During the training phase, the teaching signal ***x***(*t*) is used to perform the RLS update. Then, during the testing phase, the teaching signal is removed.

We used the parameters given in Table 4 for all FORCE experiments unless otherwise specified.

**Table 4:** FORCE network parameters

| Parameter | Value |
| --- | --- |
| $\Delta t$ | 0.04ms |
| $N$ | 1000 |
| $M$ | 500 |
| $\tau_m$ | 10ms |
| $\tau_s$ | 20ms |
| $U_{th}$ | -40mV |
| $U_0$ | 0mV |
| $U_r$ | -65mV |
| $t_{ref}$ | 2ms |
| $Q$ | 10 |
| $G$ | 0.04 |
| $A$ | 80 |

The period *T* was chosen to be equal to length of the teaching signal ***x***[*t*]. The membrane potential were randomly initialised following a uniform distribution 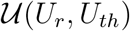. Vectors ***η*** and ***μ*** are randomly drawn from 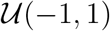. The static weights ***v***^0^ are drawn from a normal distribution 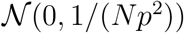, then these weights are set to 0 with probability *p*= 0.1. All other variables are initialised to zero unless otherwise specified.

##### S1. Heterogeneity on other spiking neuron hyperparameters

**Table 5:** Performance comparison among different RSNN configurations on SHD testing set. The configuration for *τ_m_* and *τ_s_* was used as specified in the top row. Initialisation and training schemes were applied only for the parameter in its corresponding column

| | **HomInit-StdTr** $\tau_m$ and $\tau_s$ | | | **HetInit-HetTr** $\tau_m$ and $\tau_s$ | | |
| --- | --- | --- | --- | --- | --- | --- |
| | $U_{th}$ | $U_0$ | $U_r$ | $U_{th}$ | $U_0$ | $U_r$ |
| **HomInit-StdTr** | $72.8 \pm 3.4$ | $72.8 \pm 3.4$ | $72.8 \pm 3.4$ | $78.9 \pm 2.0$ | $78.9 \pm 2.0$ | $78.9 \pm 2.0$ |
| **HetInit-StdTr** | $71.1 \pm 2.3$ | $69.9 \pm 2.3$ | $70.8 \pm 3.3$ | $79.2 \pm 2.9$ | $74.8 \pm 4.9$ | $79.1 \pm 2.7$ |
| **HomInit-HetTr** | $73.7 \pm 2.8$ | $71.6 \pm 3.1$ | $70.5 \pm 3.7$ | $79.1 \pm 1.8$ | $76.1 \pm 3.3$ | $76.4 \pm 2.7$ |
| **HetInit-HetTr** | $73.2 \pm 2.8$ | $72.0 \pm 2.6$ | $69.4 \pm 3.8$ | $79.2 \pm 2.8$ | $74.9 \pm 6.6$ | $75.4 \pm 6.6$ |

##### S2. Membrane time constant distribution consistency across trials

**Figure 4:**
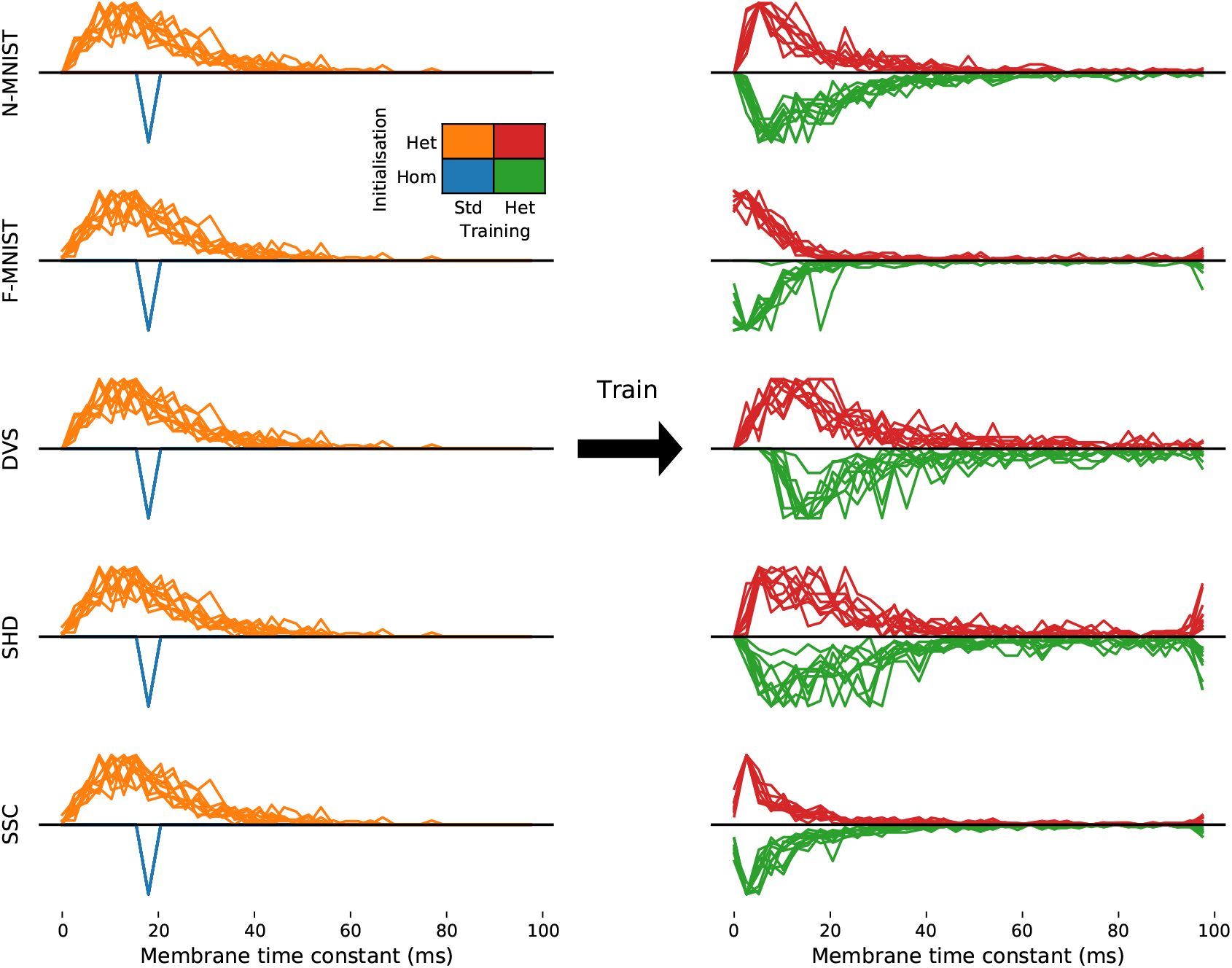
Breakdown of membrane time constant distribution for each trial for consistency checking on each dataset

##### S3. Synaptic time constant distribution consistency across trials

**Figure 5:**
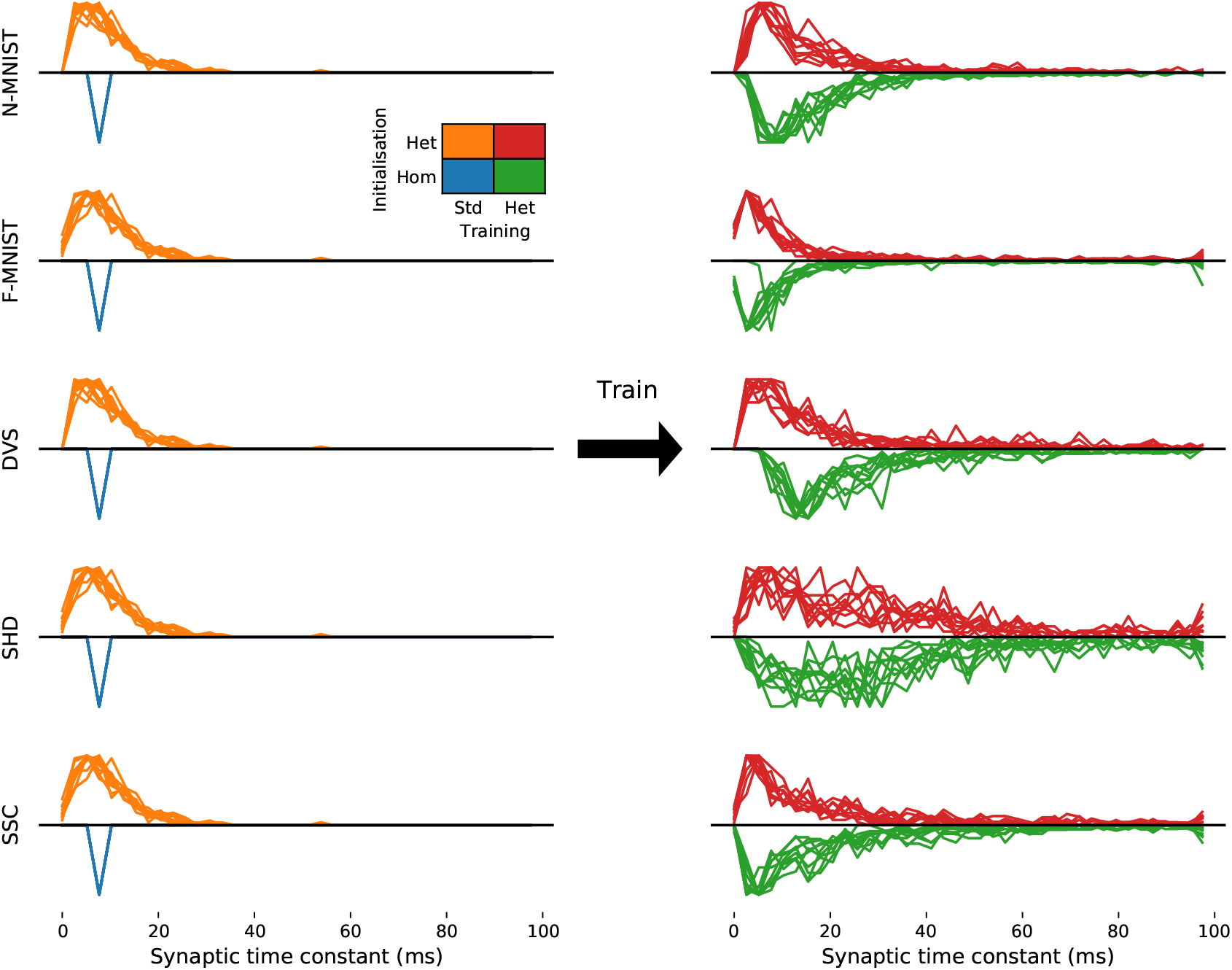
Breakdown of synaptic time constant distribution for each trial for consistency checking on each dataset

##### S4. Confusion matrices DVS-gesture dataset

The classes correspond to

1. Hand clapping
2. Right hand wave
3. Left hand wave
4. Right arm clockwise
5. Right arm counter-clockwise
6. Left arm clockwise
7. Left arm counter-clockwise
8. Arm roll
9. Air drum
10. Air guitar

**Figure 6:**
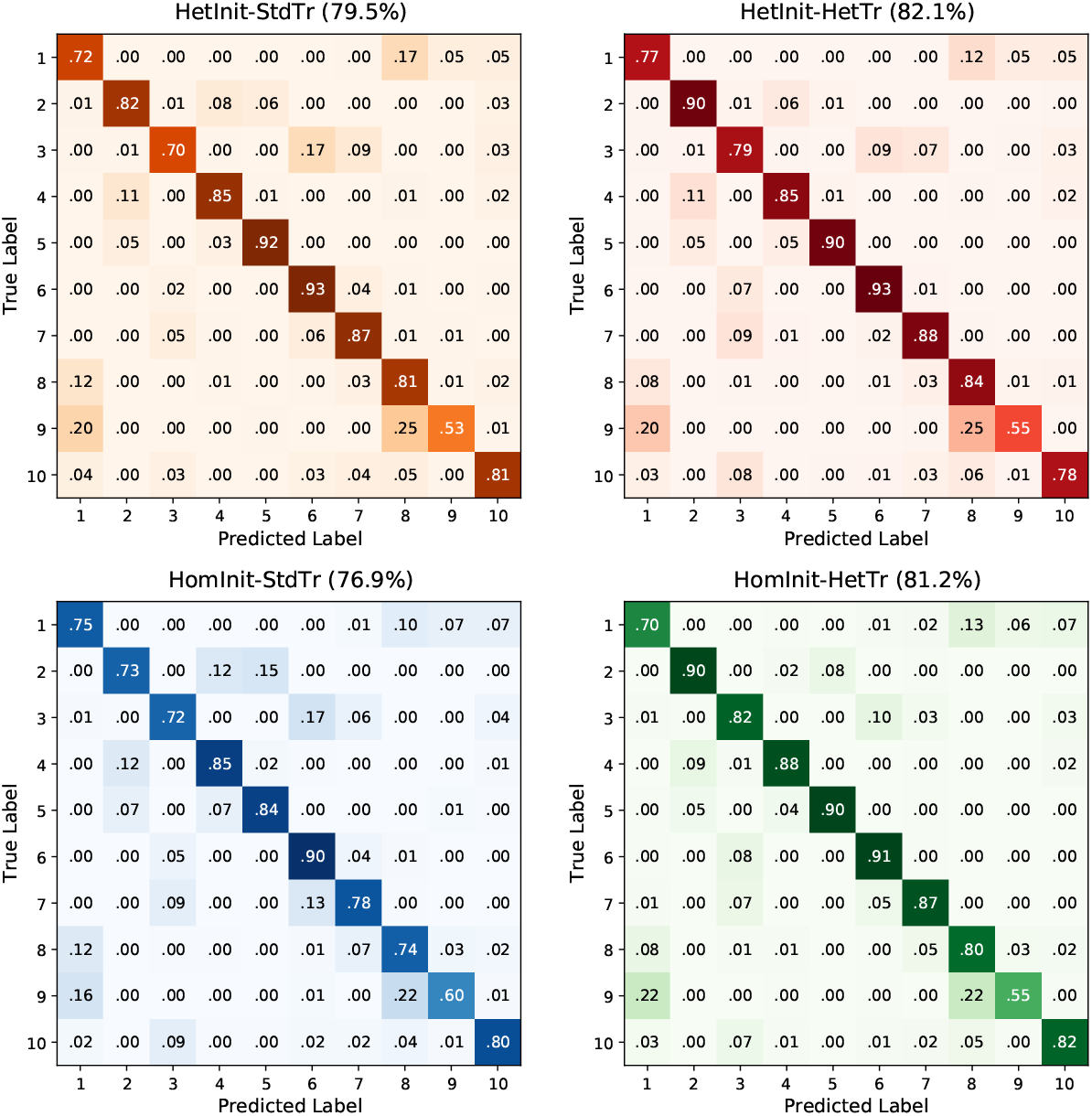
Full confusion matrix DVS dataset for each configuration

##### S5. Grid search of parameters

**Figure 7:**
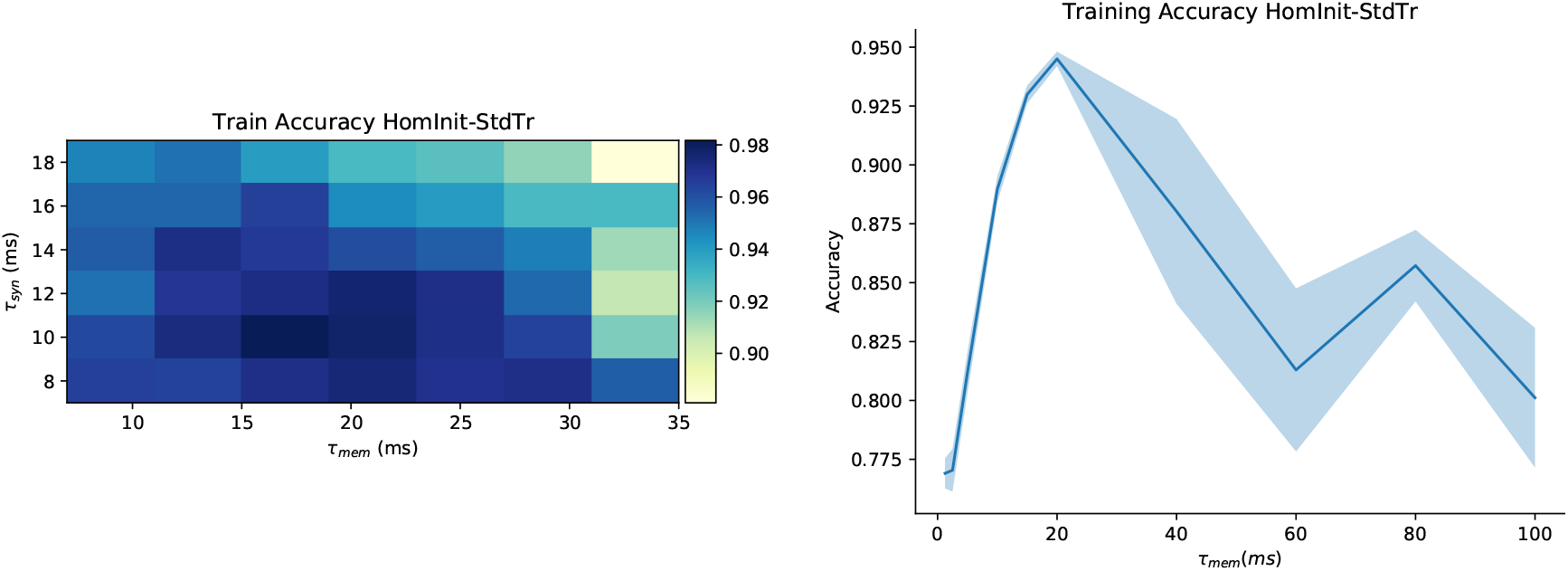
**A.** Grid search of of optimal membrane and synaptic time constants on the SHD dataset for a single trial **B.** Grid search of of optimal membrane time constant for the SHD dataset given synaptic time constant is 10ms for 10 trials. Result *τ_syn_* = 10*ms, τ_mem_* = 20*ms* which is the same as those time constants used in [14]

##### S6. All trials Fashion-MNIST

**Figure 8:**
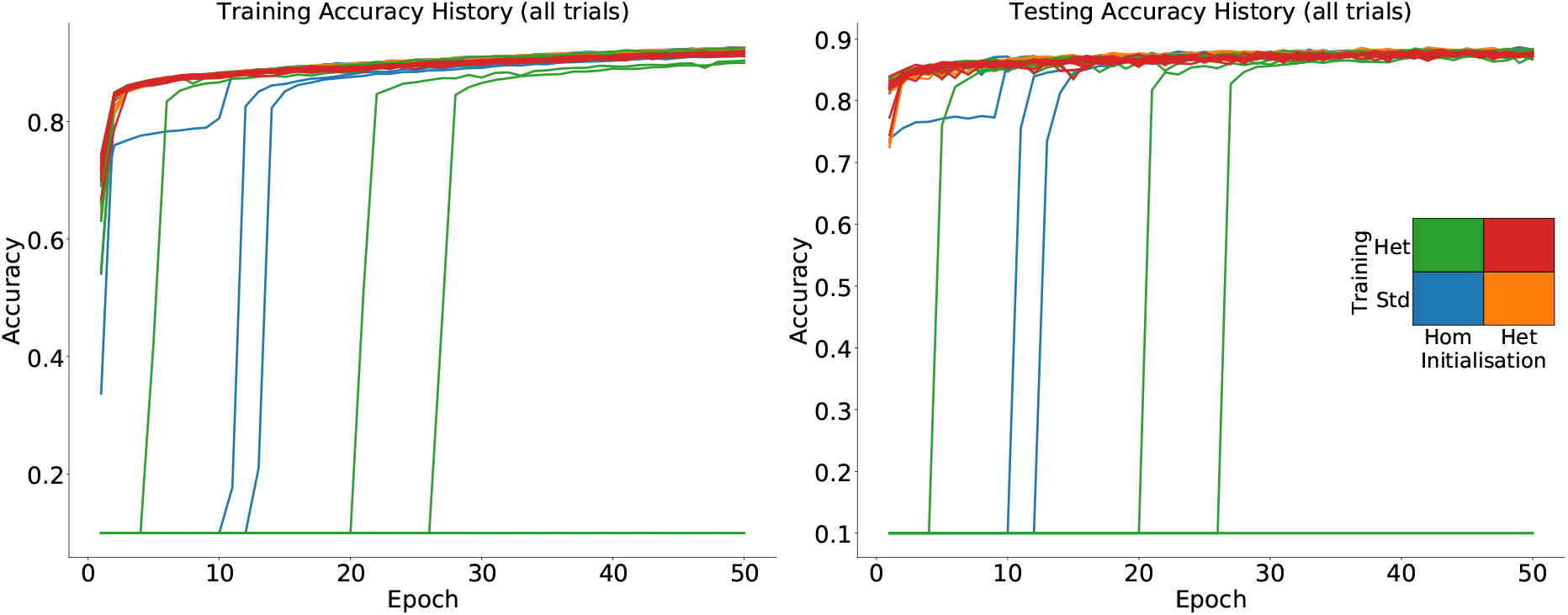
All individual trials under all four configurations (total of 40). Note how a heterogeneous initialisation results in faster convergence consistently.

##### S7. Robustness under other time constant distributions

**Figure 9:**
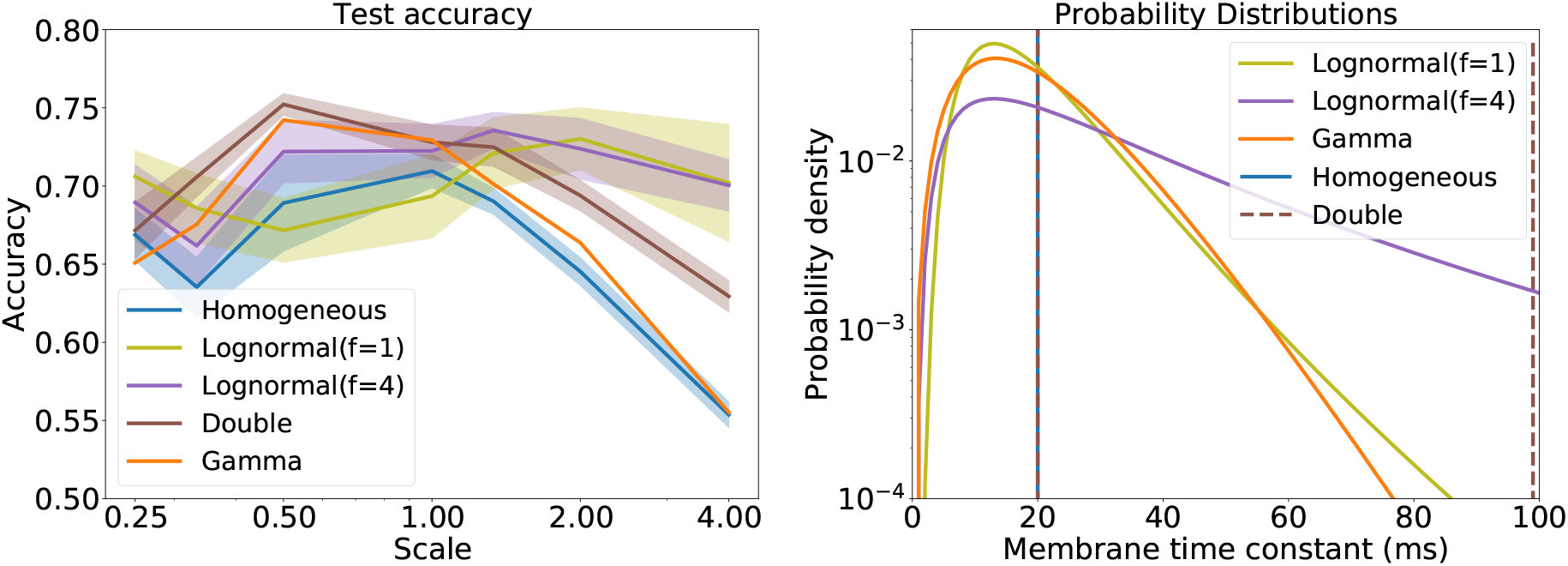
**Left**: Test accuracy with standard training and different heterogeneous initialisations for both membrane and synaptic time constants **Right**: Probability distributions for the membrane time constants (Homogeneous, Lognormal, Double (like homogeneous but 5% of the membrane and synaptic time constants are 100*ms*) and Gamma)

##### S8. Visualisation of temporal structure of DVS128 gesture dataset samples

**Figure 10:**
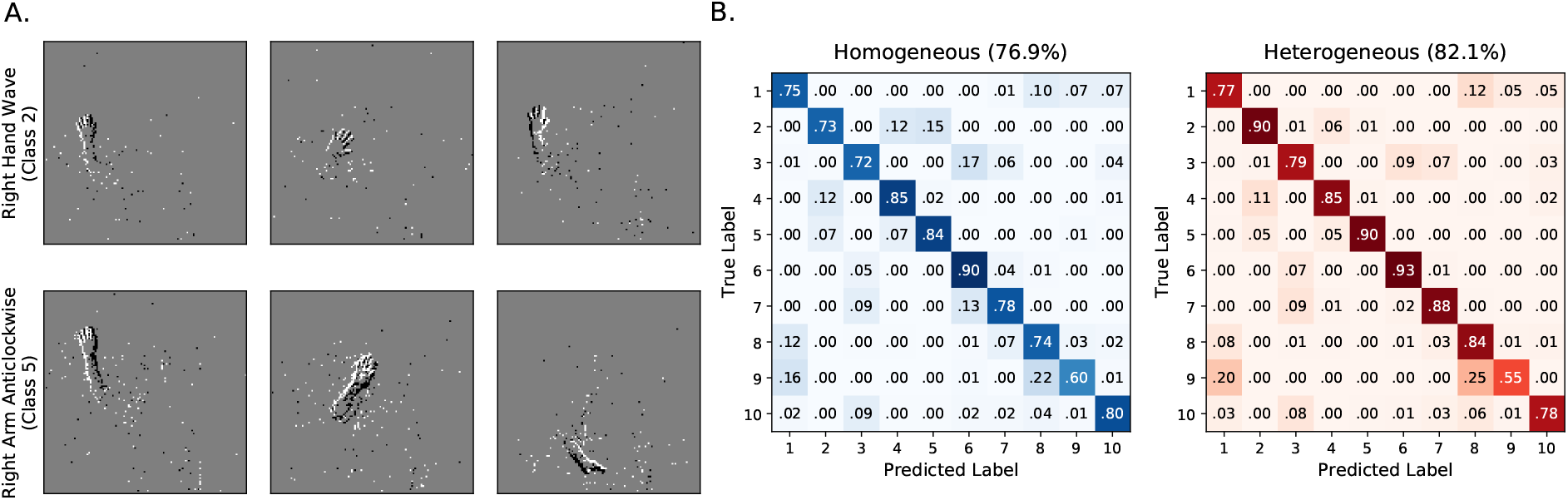
**A.** Visualization of two samples of the DVS128 gesture dataset. Each frame spans a 5ms window with 200ms between frames. To distinguish these gestures we need to integrate over tens of milliseconds. **B.** Confusion matrices of the DVS128 gesture dataset under the fully homogeneous (left) and fully heterogeneous (right) configurations. Class 2 (*Right hand wave*) is often incorrectly classified as 4 (*Right hand clockwise*) and 5 (*Right hand counter clockwise*) under the homogeneous configuration but not under the heterogeneous one.

## Notes

### Competing Interest Statement

The authors have declared no competing interest.

### Summary of Updates

Slight improvements of some results based on further simulations.

